# Epithelial Cell Regulation of Type 2 Cytokine Release by Peripheral Blood Mononuclear Cells

**DOI:** 10.1101/2020.05.01.067694

**Authors:** Amy Southern, Aurelia Gondrand, Scott Layzell, Jennifer L Cane, Ian D Pavord, Timothy J Powell

## Abstract

**Background:** Type 2 cytokines such as IL-13 and IL-5 are important drivers of pathophysiology and exacerbation in asthma. Defining how these type 2 cytokine responses are regulated is a research priority. Epithelial cells promote type 2 responses by releasing alarmins including IL-25, IL-33 and TSLP, but much less is known about inhibitory factors.

**Methods:** IL-13 release was measured from peripheral blood mononuclear cells (PBMC) cultured with Interleukin (IL)-2 for five days. Epithelial cell lines or human bronchial epithelial cells (HBEC) isolated from healthy or asthma donors were added to these PBMC cultured with IL-2 and release of IL-13 or IL-5 measured. To characterise the mechanisms, we assessed the effect of mechanical disruption of epithelial cells, addition of the COX inhibitor indomethacin and the G-protein inhibitor pertussis toxin.

**Results:** PBMC cultured with IL-2 secreted type 2 cytokines in a cell number and time dependent manner. Epithelial cell lines inhibited IL-13 and IL-5 release after co-culture with PBMC in the presence of IL-2, directly, across a transwell and using epithelial cell supernatant. Cells or supernatant from HBEC from healthy or asthma donors also inhibited the cytokine release. Trypsin treatment of conditioned media indicated that inhibitory factor(s) are trypsin insensitive. Mechanical disruption of epithelial cells or indomethacin treatment had no effect, but pertussis toxin reduced epithelial cell inhibition of IL-2 driven type 2 cytokine release.

**Conclusion:** Epithelial cells regulate cytokine release by soluble factor(s) and this could be an important immunoregulatory function of the airway epithelium.

## Introduction

Type 2 inflammation is common amongst asthma patients and is coordinated by type 2 cytokines, such as IL-13 and IL-5 resulting in eosinophil recruitment to the airways and the development of airflow limitation [1–4]. Whether epithelial cells play a role in regulating these cytokines or cells or whether epithelial cell regulation is defective in asthma patients is not fully understood [5, 6]. Epithelial cells have been shown to promote type 2 inflammation through the release of Thymic Stromal Lymphopoietin (TSLP), IL-33 and IL-25 which in the process of antigen presentation, influence dendritic cells to differentiate T cells towards a type 2 response involving IL-4, IL-5, IL-13 and granulocyte macrophage colony stimulating factor (GM-CSF) which attract eosinophils and basophils to the lung [7]. These type 2 cells can be measured in peripheral blood and sputum/lung biopsy in higher numbers in asthma patients compared to healthy controls [8–10]. However this cascade of type 2 inflammation may become excessive if it is not regulated [11]. Epithelial cells may also play a role in inhibition of this inflammatory response [12]. Using ex-vivo culture, epithelial cells have been shown to regulate T cells and to reduce the amount of cytokine secreted by these cells in the presence of antigen [13–15]. Soluble mediators have been implicated in this epithelial cell regulation because inhibition has been shown using transwells and epithelial cell supernatant [16] but this has not been the case with all studies [17].

We wished to further investigate whether epithelial cells inhibit type 2 cytokine production and the mechanism of inhibition. We used a PBMC based protocol to investigate this process and measured release of type 2 cytokines using ELISA based methods. Epithelial cell lines inhibit IL-2 driven type 2 cytokine release from peripheral blood mononuclear cells (PBMC) and this regulation is dependent upon cell numbers. Inhibition is mediated by a soluble entity as supernatants from epithelial cells inhibit IL-2 driven IL-13 release from PBMC. Human bronchial epithelial cells (HBEC) from healthy or asthma donors and supernatants derived from these cells cause a similar degree of inhibition. Trypsin treatment of the supernatant indicates that the inhibitory moiety is non-protein in nature. Mechanical disruption of the monolayer of epithelial cells and the addition of indomethacin does not reduce the type 2 cytokine inhibition. However, the addition of pertussis toxin results in the reduction of epithelial cell driven inhibition. This regulation of type 2 cytokine release could be an important potential regulator of type 2 airway inflammation.

## Methods

### Ethical Statement

Blood donations from human volunteers were taken in accordance with the declaration of Helsinki and each donor gave informed written consent. The study was approved by Leicestershire, Nottinghamshire Rutland Research Ethics Committee, UK (08/H0406/189). Blood cones were obtained from NHS blood and transplant (NHSBT), Oxford, under ethics number T298. Donors are anonymised and clinical information is not available from the cone blood.

### Cell lines, HBEC and PBMC

Lung epithelial cell lines were A549 from Dr Luzheng Xue, originally purchased from ECACC; BEAS2B and 16HBE cell lines were from Prof Ling Pei Ho, University of Oxford. A549 were maintained in D10 (DMEM with 10% v/v FCS, 2 mM Glutamine, 100 U/mL penicillin and 100 ug/mL streptomycin (all from Sigma). BEAS2B and 16HBE were maintained in F10: Ham’s F12K medium (Gibco) supplemented similarly to D10. Human bronchial epithelial cells (HBEC) for healthy controls were purchased from Lonza and grown in complete bronchial epithelial growth media (BEGM) media (Lonza). Asthma patient samples were taken from patients undergoing research bronchoscopy or clinical treatment. Cells were then maintained in BEGM media with antibiotics in a flask for 10-14 days, removed from the flask with trypsin (Sigma) and neutralised with trypsin inhibitor (Life Technologies) and then split into a 24 well plate. Cells were used at Passage (P)2 or P3 when 90% confluent. Human peripheral blood mononuclear cells (PBMC) were separated from healthy human blood samples using Ficoll-Paque (GE healthcare) or Lymphoprep (Stem Cell) by density centrifugation. PBMC were either used fresh or stored in liquid nitrogen for later usage.

### Peripheral blood cytokine stimulation assay

Epithelial cells (10^5^ cells) or media were cultured in 96 well flat bottom plates overnight then PBMC (2×10^5^) from healthy donors were cultured at 37 °C in X-VIVO 15 media (Lonza) with 50-200 U/mL IL-2 (Peprotech). The concentration of 200 U/mL IL-2 was equivalent to 20ng/mL by weight. After 5 days supernatants were harvested and cells removed and then either used straight away or frozen for later use. For some experiments different numbers of epithelial cells were placed in the wells the day before the assay. For transwell experiments, 5×10^5^ epithelial cells were added to 24 well plates and cultured overnight. Media was replaced with fresh X-VIVO media then cells were added in transwells.1×10^6^ cells in 200 uL X-VIVO with added IL-2.

For HBEC, these were cultured in 24 well plates until confluence then media was changed to X-VIVO, and PBMC added in transwells, TC insert for 24 well plate PET Membrane Bottom Transparent Pore Size 0.4um Non-Pyrogenic Endotoxin-free (Sarstedt), on top of the HBEC cells.

### Measurement of Cytokines

IL-13 was measured using an eBioscience ELISA kit (Thermo Scientific) using the manufacturer’s instructions. IL-5 and IL-2 were measured using an R&D Duoset ELISA with optimised conditions (BioTechne). In some experiments an IL-5 development ELISA was used (Mabtech). Measured concentrations were then adjusted to pg cytokine per million PBMC at the start of the assay. The range of values when PBMC were cultured with 200 U/mL IL-2 was between 143 and 1220 pg / 10^6^ cells dependent on donor. Negative donors produced undetectable concentrations of IL-13 when cultured with 200 U/mL IL-2. Repeated measurement of one donor stimulated with 200 U/mL IL-2 resulted in release of 2352, 1099, 2012, 1642, 695 and 1129 pg IL-13 / 10^6^ cells. We present here a range of donors who all produce an experimentally robust level of IL-13 or IL-5 in our assay.

### Flow cytometry

After five days of culture with epithelial cells, or IL-2 alone, remaining PBMC were harvested and labelled for different flow cytometry markers including live dead fixable viability stain eFluor780 (Thermo eBioscience cat no. 65-0865-14). Cells were then run on a BD LSR Fortessa (Becton Dickinson) and data analysed using FlowJo 10 (FlowJo & BD).

### Supernatant size exclusion and trypsin treatment

Supernatants from epithelial cells were centrifuged at 5000g for 1 hr at 4 °C and the residual volume collected using 3 kDa molecular weight cut off 50 mL or 15 mL tube filters (Millipore). The lower volume was collected and volume measured then the residual volume from the upper level was diluted to a similar volume to the lower fraction. Trypsin from porcine pancreas (Sigma T4549) was used for overnight digestion of protein (BCA, Pierce) measured supernatant and then supernatant was added to cell cultures after trypsin inactivation with trypsin inhibitor or BSA Trypsin was added at a w/w ratio of 1:50, 1:12.5 and 1:10 with epithelial cell supernatant having a protein content of 2.3 mg/mL.

### Regulation of epithelial cell inhibition

For mechanical disruption epithelial cells were plated out in 96 well plates and then after overnight cultures monolayers were disrupted with a pipette. Monolayer disruption was confirmed by microscopy. PBMC in X-VIVO with IL-2 were added and 48 hours later monolayers were disrupted again. After 5 days supernatants were collected and cytokines measured. For indomethacin 10, 25 and 50 uM of indomethacin (Sigma) was added which spanned the IC_50_ of human COX1 and COX2 throughout the course of the 5 day assay in addition to PBMC and epithelial cells. For pertussis toxin (Abcam) 1000, 500 or 200 ng/mL was added throughout the course of the assay in addition to epithelial cells and PBMC.

### Analysis and statistical tests

Data were analysed using Graphpad prism v8 using two way ANOVA with additional parametric or non-parametric tests as appropriate. In some experiments students t-test was used with correction for multiple comparisons as appropriate.

## Results

We induced IL-13 release from PBMC using stimulation from a low dose of IL-2, as previously described by Bartemes et al., [18]. This avoided potentially overstimulating the cells with anti-CD3/CD28 or PMA/ionomycin. The doses of IL-2 were similar to those used to maintain T cell clones or lines in culture [19, 20]. PBMC were isolated from donors and cultured in the presence of titrated doses of IL-2. This showed that there was an increase in the release of IL-13 (Figure 1A) and IL-5 (Figure 1B) as the concentration of IL-2 increased. Mean pg IL-13 release per 10^6^ cells +/- SD was 4.92 +/- 8.59 when PBMC were cultured in media, 171.23 +/- 144.33 with 50 U/mL IL-2, 415.72 +/- 333.94 with 100 U/mL IL-2 and 741.53 +/- 392.79 when cultured with 200 U/mL IL-2 (all n=10). Mean pg per 10^6^ cells +/- SD IL-5 release was 2.71 +/- 2.48 when PBMC were cultured in media, 23.95 +/- 19.95 with 50 U/mL IL-2, 62.52 +/- 73.76 with 100 U/mL IL-2 and 113.84 +/- 126.54 with 200 U/mL (n=5 for each). We examined the time course of this IL-13 release and found that the amount of IL-13 increased between days 2-5 and then was similar at day 7 (Figure 1C). The secretion of IL-5 into the media by PBMC also increased with time in culture (Figure 1D).

**Figure 1.**
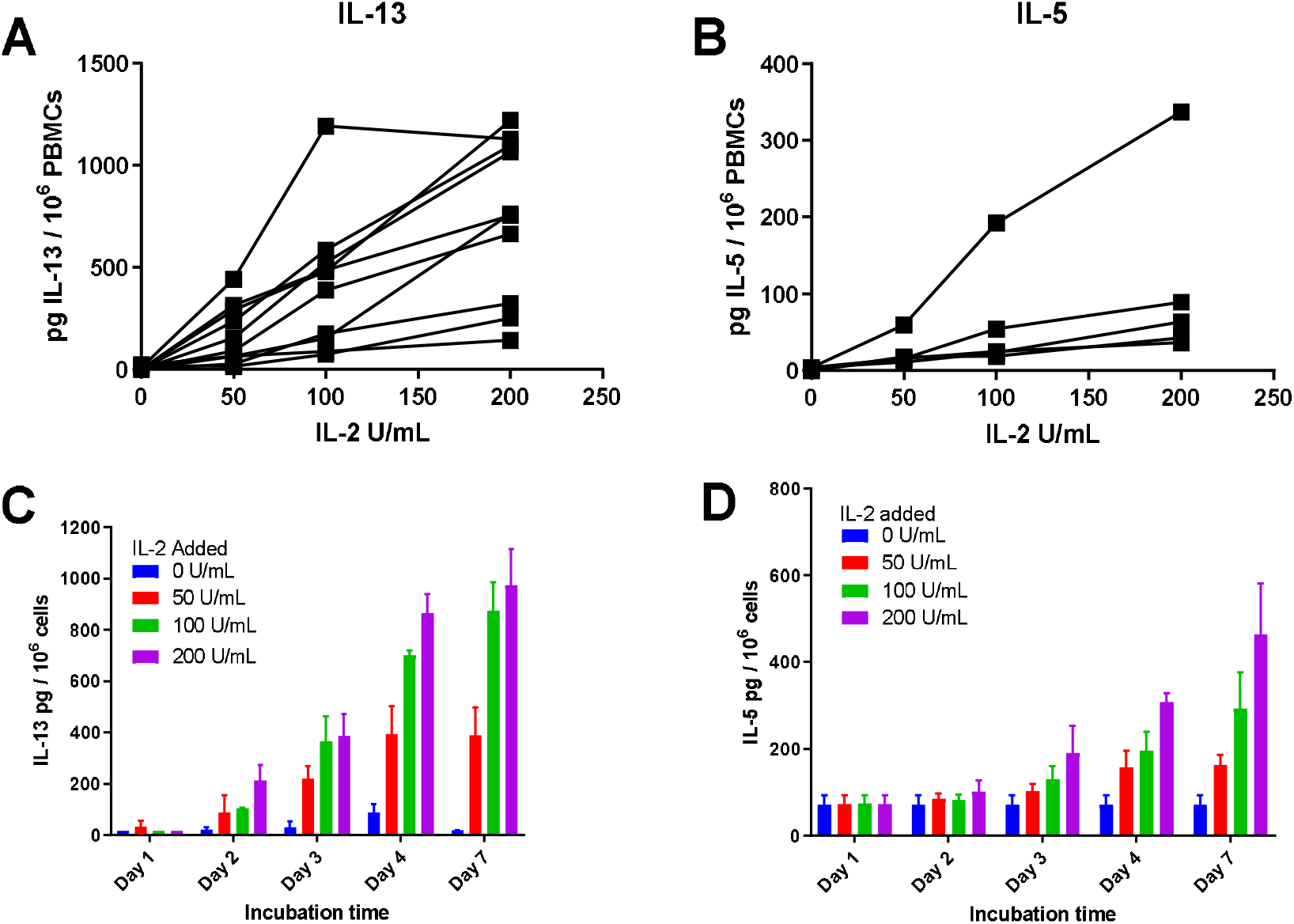
PBMC cultured with IL-2 secrete detectable levels of IL-13 or IL-5 that increase with higher concentrations of IL-2 and longer time in culture. PBMC were isolated from donors and cultured with titrated doses of IL-2 for the indicated time periods. Supernatants were collected and cells removed by centrifugation. Levels of IL-13 or IL-5 in supernatants were tested by ELISA. (A) n=10, (B) n=5 donors each tested at least once. Each dot is the mean of three experimental wells. (C, D) One representative experiment shown repeated twice. Values shown are the mean +/- SD of three wells.

When PBMC were cultured with IL-2 in direct contact with epithelial cells that had been adhered to tissue culture wells overnight, IL-13 and IL-5 release was inhibited by alveolar adenocarcinoma basal epithelial cells A549 or retro-transformed bronchial epithelial cell lines BEAS2B and 16HBE (Figure 2A-B). Mean pg per 10^6^ cells IL-13 release +/- SD when PBMC were cultured with 200 U/mL IL-2 was 513.88 +/- 397.770 (n=5) with A549 10.22 +/- 3.43 (**p=0.0036, n=3), with BEAS2B 72.18 +/- 71.86 (*p=0.0136, n=3) and 16HBE cells 32.94 +/- 32.77 (**p=0.0023, n=4). Thus culturing PBMC with IL-2 in the presence of epithelial cells led to a profound inhibition of IL-13 release. When IL-5 was measured this also showed a reduction in the presence of epithelial cells. Using 200 U/mL IL-2, the pg IL-5 per 10^6^ cells release for PBMC alone was 147.99 +/- 156.62 (n=6) and when A549 were added release was 5.94 +/- 2.09 (**p=0.0099, n=3), BEAS2B 8.17 +/- 5.46 (*p=0.0114, n=3) and 16HBE 19.34 +/- 15.13 (*p=0.0226, n=3). Therefore additions of epithelial cells to cultures of IL-2 stimulated IL-13 or IL-5 resulted in significant reduction of IL-13 or IL-5 secretion and this inhibition was most pronounced at higher concentrations of IL-2. To test its utility, the percentage reduction in IL-13 or IL-5 release was calculated for the variation in IL-2 concentration and cell type (Supp. Figure 1A, B). The 200 U / mL IL-2 was most consistent and showed the largest percentage inhibition as indicated by the IL-13 and IL-5 concentrations in Figure 2A, B. The percentage inhibition reports a negative value at low values of IL-2 because there is much less cytokine measured so a small value change can lead to a large percentage difference.

**Figure 2.**
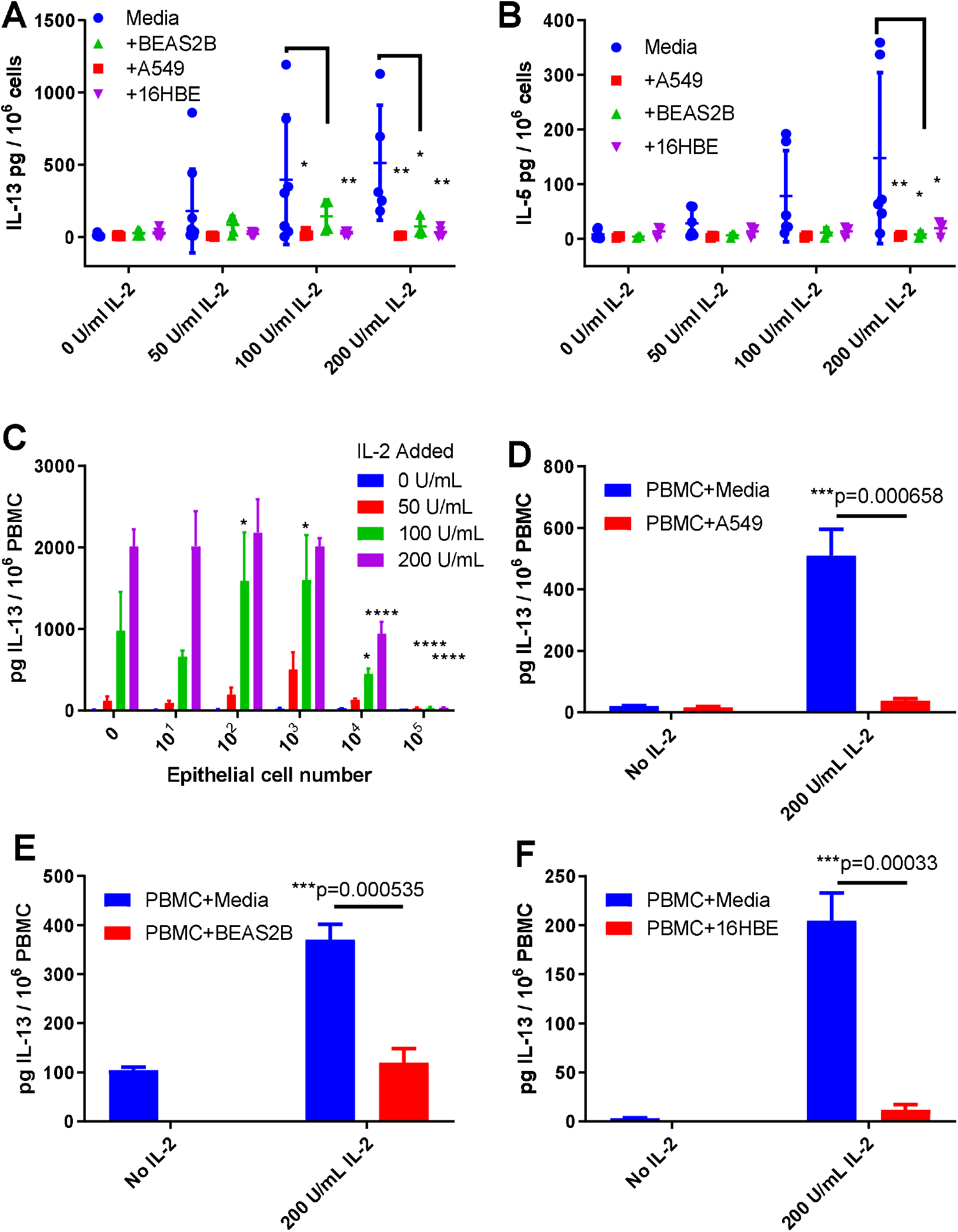
Epithelial cells cultured with IL-2 stimulated PBMC reduce resultant IL-13 release. PBMC were cultured with titrated doses of IL-2 in the absence (media) or presence of A549, BEAS2B or 16HBE cells. Cytokines in the supernatant were measured five days later. A) IL-13 release n=9, 9, 7, 5 for media (0, 50, 100, 200 U/mL respectively), n=5, 5, 4, 3 for A549 and BEAS2B, n=6, 6, 5, 4 for 16HBE, B) IL-5 release n=6 for media, n=3 for A549, BEAS2B and 16HBE. Each dot shown is a mean of three experimental wells. Statistical comparisons are from two way ANOVA followed by Tukey multiple comparison correction *p<0.05, **<0.001. C) Titrated numbers of A549 cells as shown were added to the wells the day before the PBMC were added. Values shown are mean +/- SD of three experimental wells. Two way ANOVA followed by Dunnett’s multiple comparison correction: *p<0.05, ****p=0.0001. D) A549, E) BEAS2B or F) 16HBE cells (10^6^) were added into a tissue culture well overnight then PBMC (10^6^) added into a transwell above with or without 200 U/mL IL-2 and cultured for 5 days. Unpaired multiple t tests comparison with Holm-Sidak multiple comparison correction is shown with exact p values. All experiments were repeated at least three times.

To determine whether inhibition was due to the number of epithelial cells added to the culture wells, PBMC were cultured with IL-2 and the amount of IL-13 release was measured in the presence and absence of titrated numbers of epithelial cells. When A549 cells were added from 10^1^ up to 10^5^ epithelial cells per well, inhibition increased with increasing epithelial cell numbers over 10^4^ cells (*p<0.05 100 U/mL IL-2 and ****p=0.0001 200 U/mL IL-2) and the highest inhibition was seen with 10^5^ A549 cells added (****p=0.0001) (Figure 2C). This inhibition was seen at two doses of IL-2, 100 and 200 U/ml, and to a lesser extent at 50 U/mL. Thus IL-13 inhibition was associated with increasing numbers of epithelial cells. To test whether epithelial cells were absorbing the cytokines we added IL-13 or IL-2 to confluent epithelial cells and after overnight (IL-13) or two day culture (IL-2) the remaining amount of IL-13 or IL-2 in the cultures were measured (Supp. Figure 2). There was no difference between the cytokines cultured alone or with epithelial cells indicating that the epithelial cells were not actively absorbing the cytokines. Experiments with flow cytometry at the end of the five day culture period indicated that the BEAS2B cells (or A549 not shown), did not result in an increased rate of cell death as measured by a flow cytometry live dead stain (Supp. Figure 3). In other experiments (not shown) there was no evidence to suggest that the epithelial cells enhanced the death of the PBMC.

To test whether inhibition required cell-cell contact, we measured the inhibition of the type 2 cytokine release by the PBMC cultures when the epithelial cells were separated from the PBMC by a transwell such that only soluble factors could mediate the inhibition. We found that inhibition by A549 cells of the IL-2 driven IL-13 release was still present (Figure 2D). We also found similar results with BEAS2B (Figure 2E) and 16HBE cells (Figure 2F).

To determine whether conditioned media (CM) alone from epithelial cells could directly inhibit the release of IL-13 by PBMC cultured with IL-2. CM was directly added to PBMC cultured with IL-2 and inhibited the release of IL-13 by PBMC in a concentration dependent manner: 50% v/v reduced IL-13 more than 25% v/v (Figure 3A). Comparing the groups 25% (**p=0.0018, n=15) and 50% (**p=0.001, n=17) 16HBE CM as well as 50% v/v BEAS2B CM (*p=0.0286, n=15) were statistically different by Wilcoxon matched pairs signed ranked test. We also found that inhibition by CM was not affected by trypsin treatment (Figure 3B) and was seen in fractions above and below 3kDa (Figure 3C). These experiments indicated that the inhibitory moiety may be non-peptide in nature and was not effectively fractionated by size exclusion centrifugation.

**Figure 3.**
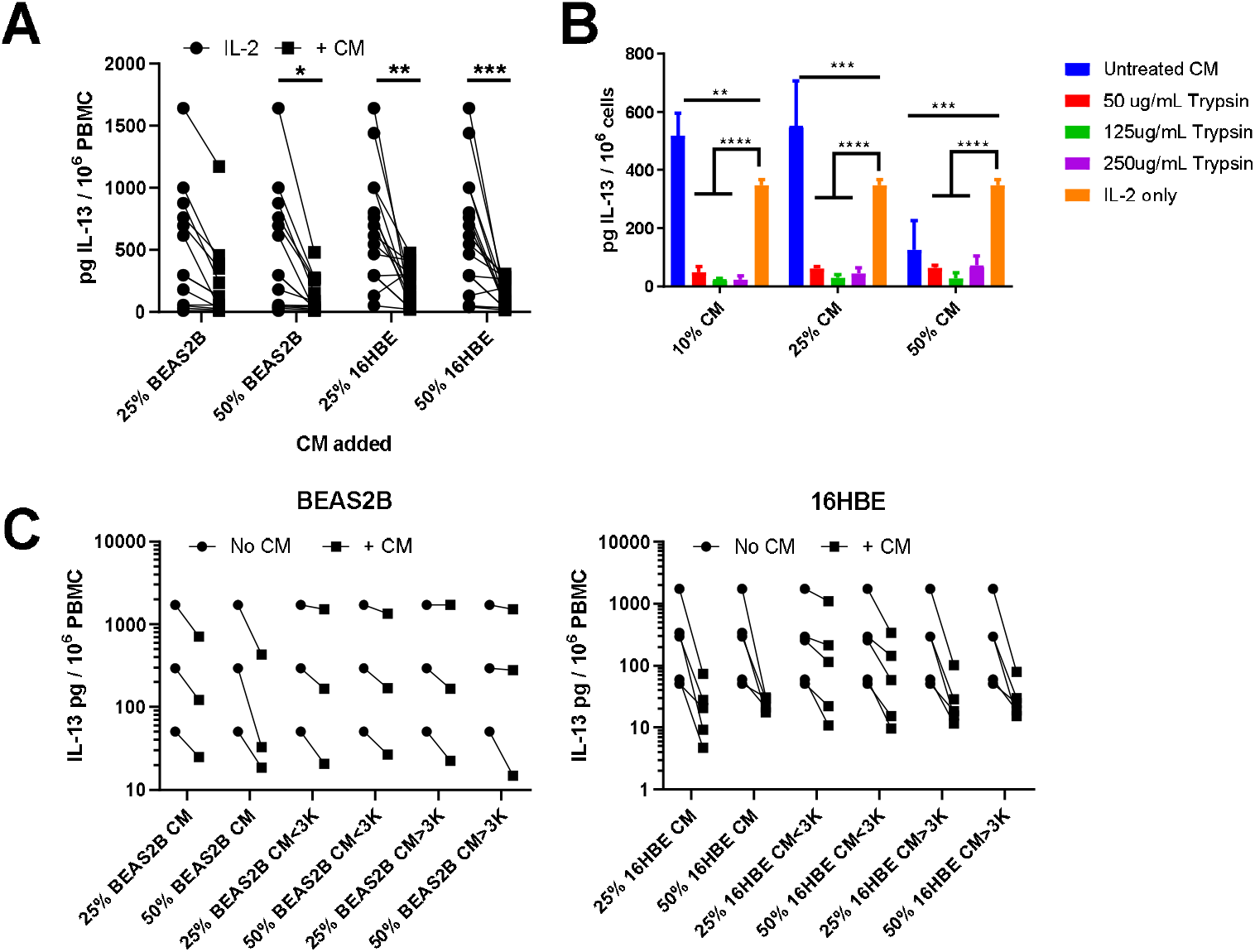
Inhibition of IL-2 driven IL-13 secretion by epithelial cell supernatant trypsin treatment and size separation. PBMC were cultured with 200 U/mL IL-2 and varying percentages of epithelial cell supernatant (CM) and then after 5 days release of IL-13 was measured. A) Increasing percentages of CM shown with IL-2 only in comparison *p<0.05 by Wilcoxon signed rank test. B) 16HBE epithelial cell supernatants were treated with trypsin overnight then trypsin quenched with and equal volume of BSA. Supernatants were then added in titrated amounts to PBMC with added IL-2 and five days later contents of IL-13 in the supernatant were measured. **p=0.0025, ***p=0.0003, ****p=0.0001 from two way ANOVA with Dunnett’s multiple comparison correction. C) CM was centrifuged through size selective filters. Titrated supernatant was then used in similar assay to A. BEAS2B n=3 and 16HBE n=5.

Then we tested whether HBEC cells were able to inhibit IL-13 release from PBMC and we found that cells derived from both healthy donors and asthma patients were able to inhibit IL-13 release from PBMC cultured with IL-2 similarly (Figure 4A). Supernatants from H.HBEC and A.HBEC cells were able to inhibit IL-2 driven IL-13 release from PBMC similarly (Figure 4B, C) and size exclusion centrifugation using a 3 kDa cut off filter did not identify a more-inhibitory fraction (Figure 4D). These data indicated that the inhibitory activity in supernatant from HBEC cells had functions in common with alveolar or bronchial cell lines shown in previous figures and that inhibition could be demonstrated across a transwell.

**Figure 4.**
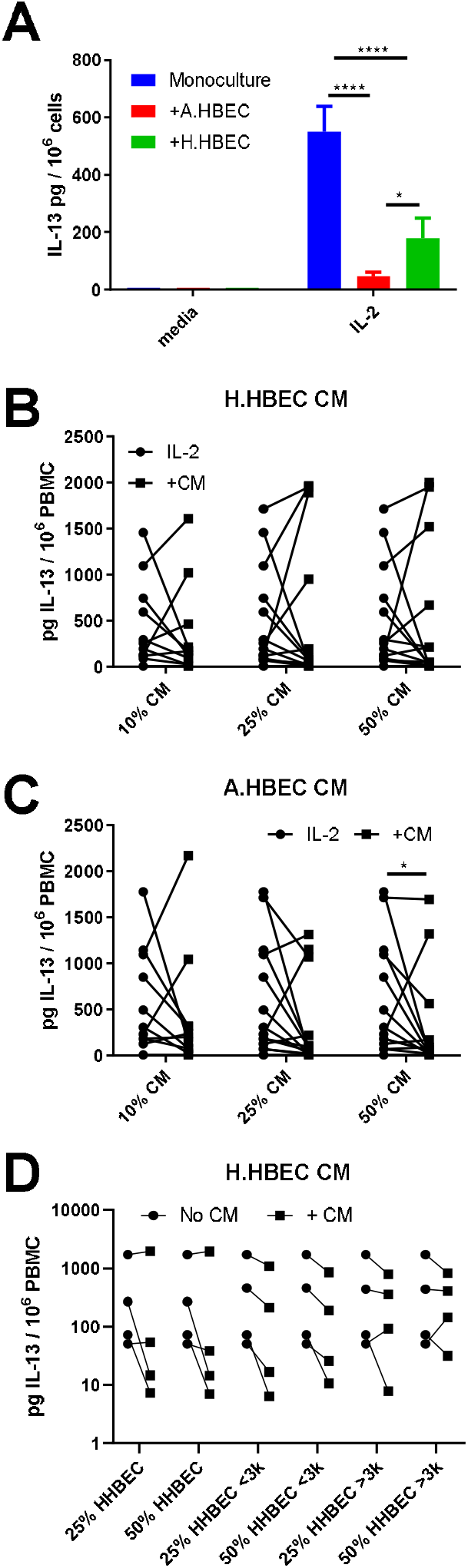
HBEC cells inhibit the release of IL-13 by PBMC but CM inhibition is not different between healthy and asthmatic epithelial cells and is not affected by CM size separation. A) PBMC were cultured with 200 U/mL IL-2 in the presence of HBEC across transwells for five days and then supernatant collected and IL13 measured. *p<0.05, ****p<0.0001 by two way ANOVA followed by Tukey multiple comparison test. B, C) CM was collected from HBEC cells as media was changed every two days. Supernatants were clarified by centrifugation then used in titrated doses in culture experiments with PBMC as above. B) Healthy HBEC CM (H.HBEC) n=14, C) Asthma HBEC CM (A.HBEC) n=14. *p<0.05 by Wilcoxon signed rank test. D) H.HBEC supernatants were size separated using centrifugal filtration devices and the resulting supernatants tested at one concentration 50% v/v and 25% v/v with IL-2 treated PBMC and IL-13 levels measured 5 days later. H.HBEC n=4.

The mechanism of suppression by the epithelial cells has not been clearly defined and since epithelial cell damage, prostaglandins and G-protein mediated pathways have been implicated in asthma pathogenesis we wished to test whether these pathways were involved with the epithelial cell inhibition of type 2 responses. Mechanical disruption, using a pipette tip, on day 0 and day 2 of culture, had no effect on the inhibition of IL-13 secretion by the epithelial cells (Figure 5A). When indomethacin was added at three different concentrations that span the IC_50_ of human COX1 and COX2 this had no effect on the epithelial cell inhibition of IL-13 release (Figure 5B). In contrast addition of pertussis toxin at a concentration of 1ug/mL did reduce the amount of epithelial cell driven IL-13 inhibition but lower concentrations of pertussis toxin had no effect (Figure 5C).

**Figure 5.**
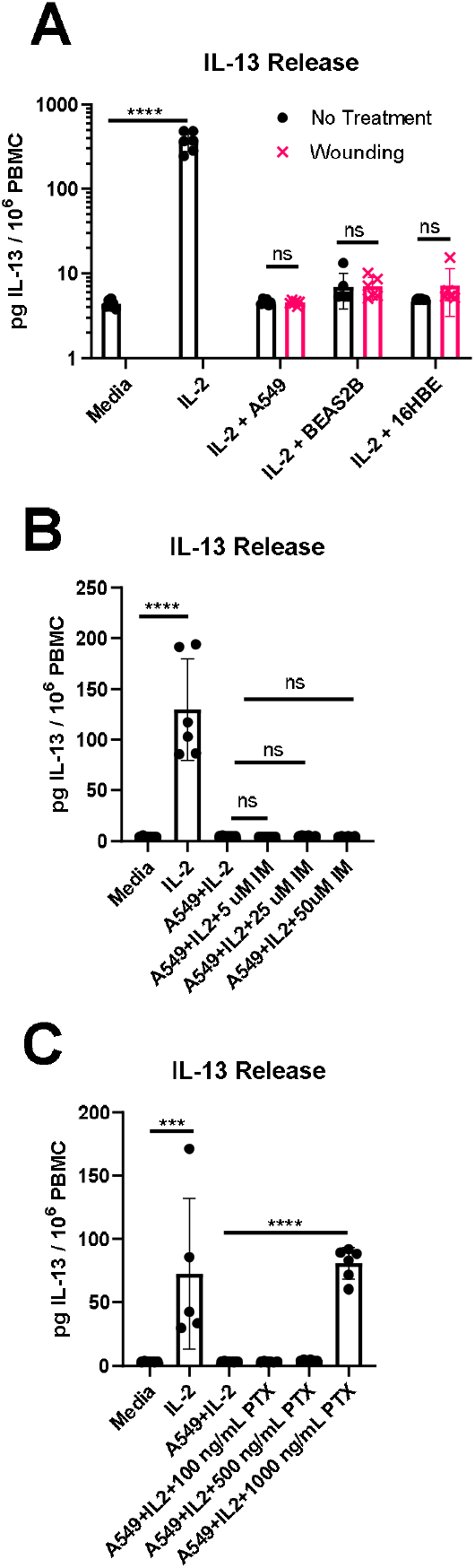
Pertussis toxin, but not indomethacin or mechanical disruption of the monolayer can reduce the amount of inhibition of type 2 cytokine release. PBMC were cultured with 200 U/mL IL-2 in the presence or absence of epithelial cells and five days later contents of IL-13 or IL-5 in the supernatant measured. Epithelial cell regulation of PBMC type 2 cytokine release was disrupted by: A) Mechanical disruption at day 0 and day 2, ****p<0.0001 by one way ANOVA with multiple comparison and Dunnett correction, ns multiple t tests with Dunnett correction. B) Addition of indomethacin ns by one way ANOVA with Dunnett correction or C) Addition of pertussis toxin. ***p=0.0001, ****p<0.0001 by one way ANOVA with Dunnett multiple comparison correction. Data shown are from a representative experiment repeated at least four times.

## Discussion

Here we observed that epithelial cells were able to inhibit the IL-2 driven secretion of IL-13 or IL-5 by PBMC. Inhibition was not dependent on cell-cell contact and was demonstrated using epithelial cell conditioned media. PBMC are a standard mixed population of immune cells that can be easily obtained from patient samples or healthy donors without enrichment of specific cell populations which gives the advantage of not selecting for nor excluding any particular type of cells that secrete type 2 cytokines. Since there is similar inhibition of IL-13 or IL-5 release as that seen with purified CD4 cells [16], it is likely that memory type 2 T cells within the PBMC are producing the IL-13 / IL-5 since these express the IL-2 receptor and have been shown to respond to IL-2 [21, 22]. It is also possible that other cells could be producing the type 2 cytokines that also express the IL-2R either directly or via the IL-2 signalling of the T cells [18]. We have cultured fresh and frozen cells from different donors and find that PBMC from most donors cultured with IL-2 secrete type 2 cytokines but this is not the case with all donors, potentially reflecting different memory cell populations, different kinds of infectious or allergen exposure, or differential sensitivity to IL-2 [21].

Our data contrast with findings from Deppong et al., [17] and Wang et al., [14] who characterised this epithelial cell regulation as mainly cell contact dependent and may constitute an additional regulatory pathway. Others have seen inhibition by epithelial cell media [13, 16] and the difference may reflect different culture conditions or different treatment of epithelial cell supernatant. We have tested 48 hour culture supernatant from different densities of HBEC cells and found that the majority of supernatants inhibit IL-2 driven IL-13 release (Figure 4B, C). However some CM does not inhibit and this could be due to an absence of cross talk between epithelial cells and PBMC because only epithelial cell CM is present [14, 17, 23].

Our data are consistent with Schwarze et al., that asthma HBEC supernatant are not less inhibitory than supernatant from healthy HBEC particularly in cytokine release [16]. The inter-donor variability we observe, in which some HBEC supernatants may be less inhibitory or even stimulatory may be apparent because there are fewer cells in the HBEC cultures compared to cell line cultures. When epithelial cell input numbers are controlled it can be shown that lower cell numbers resulted in less inhibition (Figure 2). Others have shown that bronchial biopsies and cultured epithelial cells from asthma donors are different to those from healthy controls, particularly in barrier function [24]. In addition epithelial cells derived from asthma donors may be less effective at controlling virus infection [25], cellular bounce [5], controlling eosinophil progenitors [26] or tight junction formation [27]. Airway liquid interface (ALI) cultures of epithelial cells from asthma patients were less effective than from healthy controls at reducing the release of histamine from mast cells [23] or division of CD4+ T cells [16]. However others have not shown a difference in viral control between epithelial cells derived from healthy donors and asthma patients [28, 29]. Cells cultured from patients may represent a minority population of healthy cells whilst damaged cells from asthma patients may survive less well in vitro. Thus after prolonged culture, cells from asthma patients may be more similar to those in health, reducing the phenotypic difference between healthy and asthma donors [29]. We do not have a huge cohort of healthy versus asthma donors’ epithelial cells and so there could be more of a difference if we collected more samples. In addition we didn’t see an increased rate of cell death in the PBMC cultured in the presence of epithelial cells which agrees with other investigators [23].

What is likely to be the nature of the inhibitory mediator(s)? Factors such as resolvins are candidates since these are able to inhibit T cell responses, both in bulk culture of PBMC and using purified CD4 cells [30]. However it could be a lipid mediator such as PGE_2_ [31] or PGI_2_ [32]. The non-protein nature of the inhibitory effect indicate that these mediators could be involved. However since the addition of indomethacin does not reduce the inhibition of the type 2 cytokine release this seems unlikely to be a specific mechanism but other more specific inhibitors for different classes of prostaglandin inhibitors may be useful to determine this. Pertussis toxin was able to reduce the inhibition of the IL-2 driven type 2 cytokine release and others have suggested this indicates that other prostaglandins are involved [23]. The concentration used was similar to Martin et al., [23] and in lymphocyte homing inhibition assays [33] which are quite high concentrations when compared to those used in purified protein assays [34, 35]. There are a plethora of G protein coupled responses that could be indicated by this experiment and investigation of these is underway. The other mechanism that could be involved is that cellular damage of the epithelial barrier may lead to a lack of an inhibitory signal. We tested this using mechanical disruption of the monolayers and did not see an effect. This may reflect that the mechanically disrupted cells are restored during the time of the assay and continue to inhibit the response of IL-2 driven type 2 cytokines.

In summary, we have shown that both cells and supernatants from epithelial cell lines and HBEC inhibits the release of T cell cytokines but supernatants from submerged HBEC may not inhibit type 2 cytokines differentially between asthma and healthy donors, although a larger group of asthma donors and healthy controls would provide greater power to detect smaller effects. This pathway of epithelial cell driven inhibition is reproducible across different laboratories and further work is necessary to identify the factor(s) responsible which our data suggest is a non-protein molecule released by epithelial cells that is able to inhibit type 2 cytokine release in vitro. Further elucidation of this regulatory pathway could lead to identification of novel therapeutic targets.

## Acknowledgements

The authors acknowledge the support of members of the Respiratory Medicine Unit for discussion and help in the laboratory and for assistance in obtaining clinical samples.

---

**Supplementary Figure 1.**
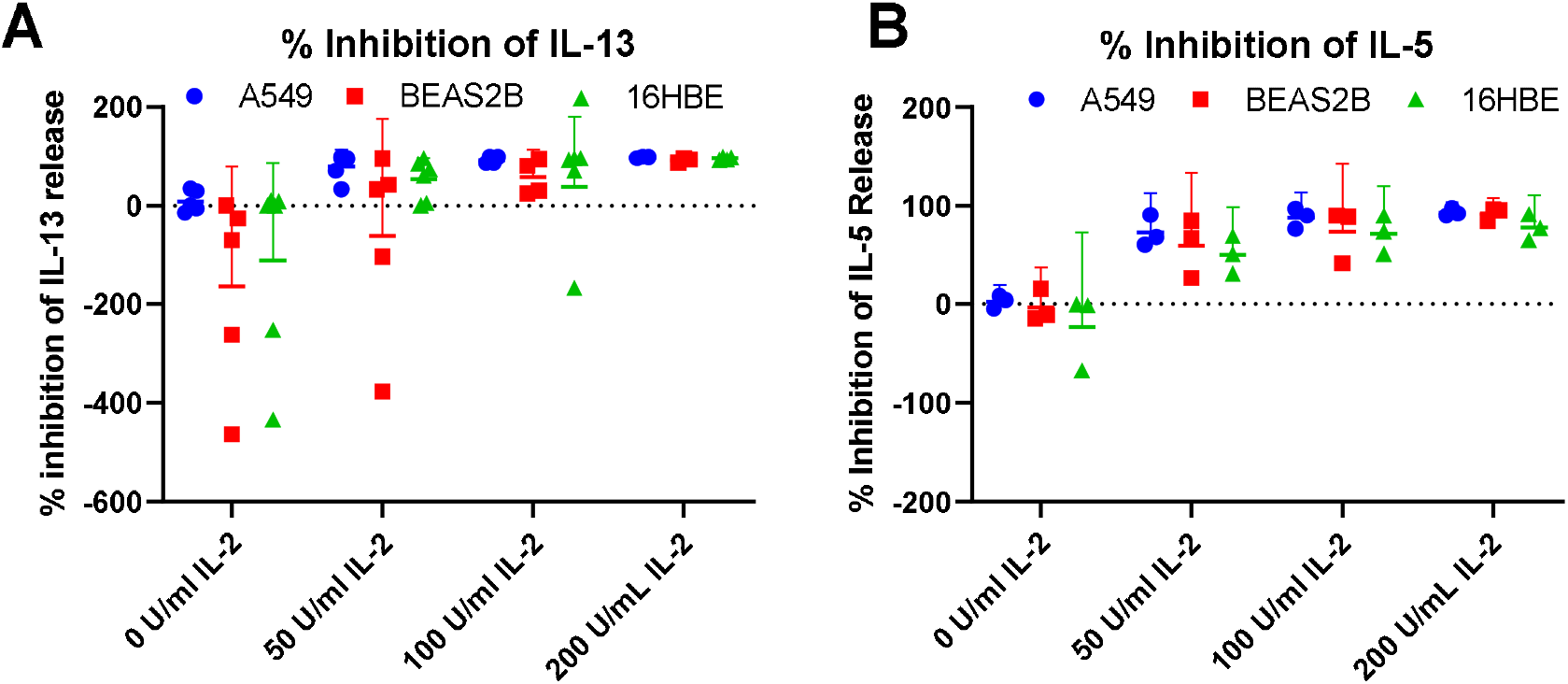
Calculation of percentage inhibition of Il-13 and IL-5 release with PBMC cultured with titrated doses of IL-2 and A549, BEAS2B or 16HBE cells. A) IL-13 or B) IL-5 concentrations were measured when PBMC (2×10^5^ cells) were cultured with titrated doses of IL-2 in the presence or absence of 10^5^ A549, BEAS2B or 16HBE cells placed in plates the previous day. Five days later cytokines were measured and percentage release calculated by dividing the (amount in the absence of epithelial cells – amount in the presence of epithelial cells) by the amount in the absence of epithelial cells multiplied by 100. Values shown are means +/- 95% CI. A) A549, BEAS2B n = 5, 5, 4, 4 and BEAS2B n=6, 6, 5, 4 for IL-2 at 0, 50, 100, 200 U/mL respectively.

**Supplementary Figure 2.**
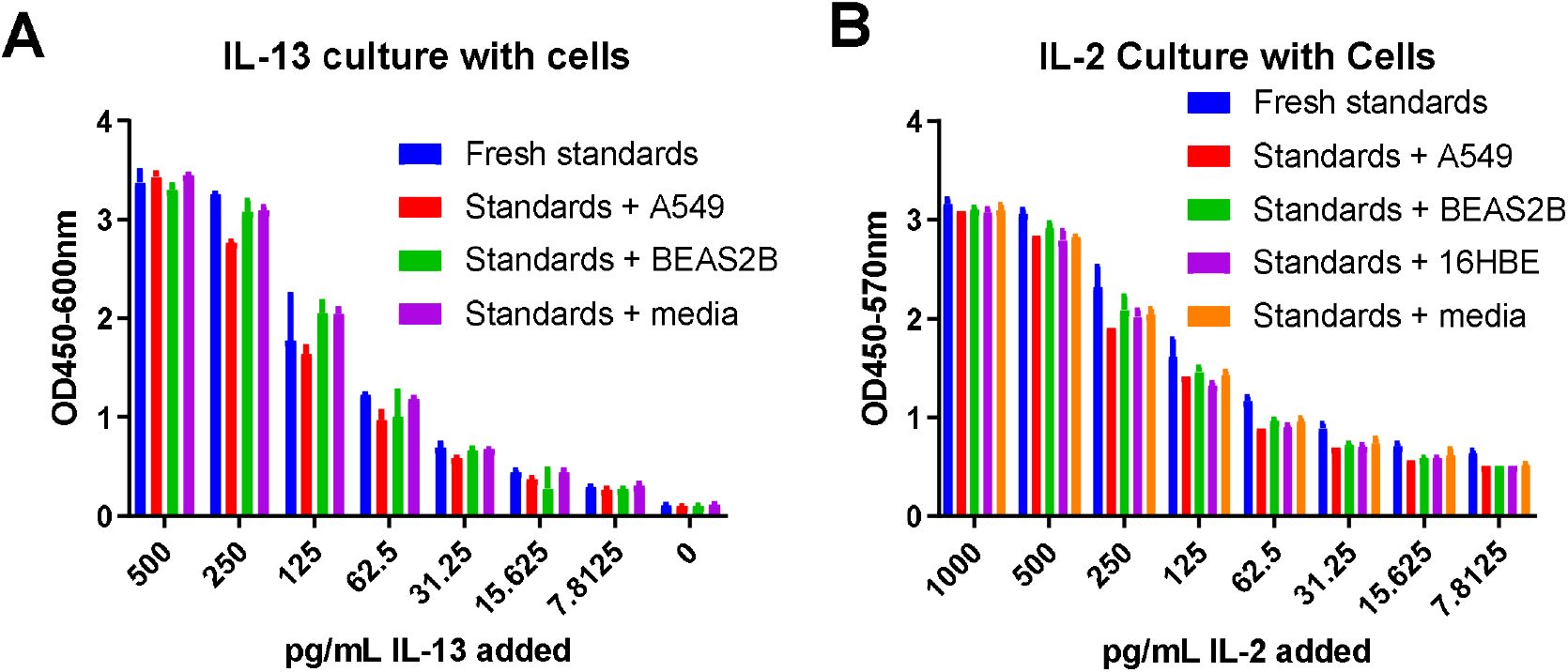
Culture of IL-13 or IL-2 ELISA standards with epithelial cells does not result in uptake of cytokine. Epithelial cells were seeded into 96 well plates overnight (1×10^5^ / well) then titrated doses of (a) IL-13 and (B) IL-2 were added. After (A) overnight and (B) 48 hours supernatants were harvested and the amount of cytokine remaining measured by ELISA. One representative experiment repeated twice is shown.

**Supplementary Figure 3.**
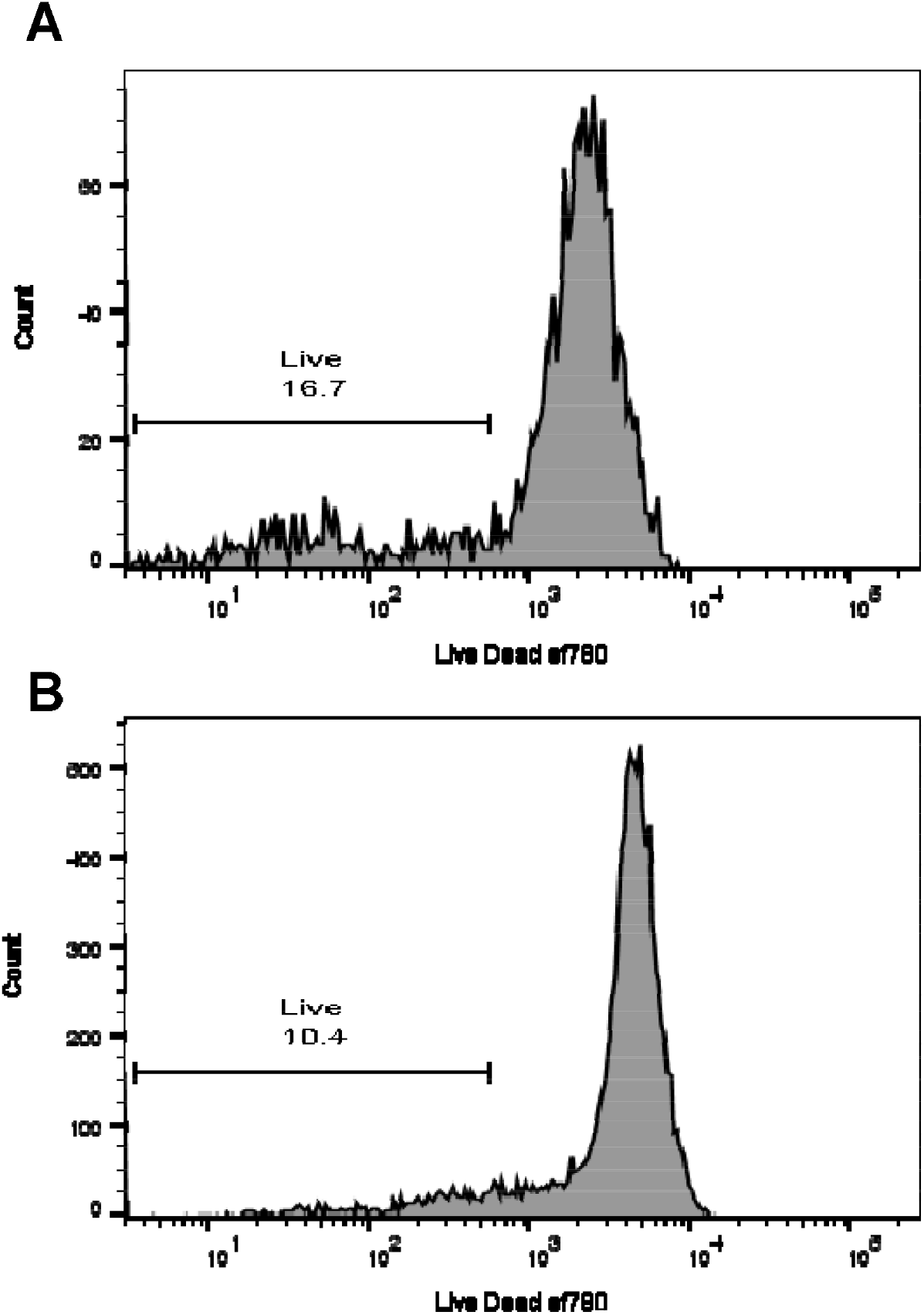
Culture of PBMC with epithelial cells does not result in increased cell death. PBMC were cultured with 200 U/mL IL-2 in the presence of absence of BEAS2B cells for five days and then labelled for flow cytometry including live dead ef780. Profiles shown are gated for singlets in a lymphocyte size / scatter gate labelled for Live / Dead efluor 780. A) PBMC cultured with BEAS2B cells and 200 U/mL IL-2 B) PBMC cultured with 200 U/mL IL-2. Gates shown are percentage live cells.

## Abbreviations

HBEC: human bronchial epithelial cells
A.HBEC: HBEC from asthma patients
H.HBEC: HBEC from healthy controls
PBMC: peripheral blood mononuclear cells
IL: interleukin
TSLP: thymic stromal lymphopoetin

